# A neurocomputational model of observation-based decision making with a focus on trust

**DOI:** 10.64898/2026.03.24.713845

**Authors:** Azadeh Hassannejad Nazir, Jeanette Hellgren Kotaleski, Hans Liljenström

## Abstract

As social beings, humans make decisions partly based on social interaction. Observing the behavior of others can lead to learning from and about them, potentially increasing trust and prompting trust-based behavioral changes. Observation-based decision making involves different neural structures. The orbitofrontal cortex (OFC) and lateral prefrontal cortex (LPFC) are known as neural structures mainly involved in processing emotional and cognitive decision values, respectively, while the anterior cingulate cortex (ACC) plays a pivotal role as a social hub, integrating the afferent expectancy signals from OFC and LPFC.

This paper presents a neurocomputational model of the interplay between observational learning and trust, as well as their role in individual decision-making. Our model elucidates and predicts the emotional and rational behavioral changes of an individual influenced by observing the action-outcome association of an alleged expert. We have modeled the neurodynamics of three cortical structures (OFC, LPFC, and ACC) and their interactions, where the neural oscillatory properties, modeled with Dynamic Bayesian Probability, represent the observer’s attitude towards the expert and the decision options. As an example of an everyday behavioral situation related to climate change, we use the choice of transportation between home and work. The EEG-like simulation outputs from our model represent the presumed brain activity of an individual making such a choice, assuming the decision-maker is exposed to social information.

## 1. Introduction

Observational learning plays an important role in many forms of human decision making, particularly in situations where individuals rely on the experiences and outcomes of others to guide their own choices. In such contexts, trust in the observed agent and the evaluation of observed outcomes influence whether individuals adopt or reject a particular strategy. Understanding these processes requires integrating insights from both social processes and the neural systems that support learning and valuation. This study therefore considers observational learning and trust-based decision making from both social and neural viewpoints, providing a framework that links behavioral processes with their possible neural substrates. Reviewing these perspectives provides a basis for introducing a neurocomputational framework that links these two levels of analysis through a computational model.

### 1.1. Background

To provide the necessary context for the present study, we first review relevant perspectives on observational learning and trust-based decision making at both the social and neural levels.

#### 1.1.1. Social-based decision making

Human decisions are influenced by information from various sources in the environment. Decision-making and learning are two concepts strongly linked together, with each serving as an inseparable basis for the other (Gold & Shadlen, 2007; Lee et al., 2012). The goal of human decision-making could be considered maximizing its utility (Doya, 2008; Fobbs & Mizumori, 2014), where different types of contextual learning (i.e. individual and social context) contribute to the knowledge base for reaching a decision (Gariepy et al., 2014; Lee & Harris, 2013).

While an individual’s own experiences may be the primary reason behind his/her learning, learning from others, i.e., social learning, also plays an important role (Campbell-Meiklejohn et al., 2010). According to Bandura (1977), social learning is based on observation, imitation, and cognitive modeling. In this context, individual learning by observing the behavior of others is known as *observational learning* (Ramsey et al., 2021). Bandura (1961; 1971) demonstrates different forms of observational learning. Encoding the observed behavior through *copying*, regardless of the potential consequence, is one form of observational learning. Another form is *observational associative learning*, when the behavioral learning could result from associating an observed action with its outcome. Allowing for conditional association between the observed action and the outcome, observational learning is the corollary of the generation of *prediction error* (PE) signals. Two forms of vicarious prediction error signals, observational *action* prediction error and observational *outcome* prediction error, i.e., the difference between the anticipated and actual value of the others’ behaviors and their outcomes, can explain human observational learning. The sign and magnitude of these signals determine the motivations for engaging in (i.e. rewarding) or withdrawing from (i.e. aversive) a specific behavior (Apps et al., 2015; Bandura, 1971; Burke et al., 2010; Chang et al., 2011; Monfardini et al., 2013; Selbing et al., 2014; de Visser et al, 2018; Kang et al., 2021; Pan et al., 2023).

The concept of associative learning is strongly related to the concept of goal-directed behavior. According to Dickinson and Balleine (1994; 2002), goal-directed behavior is based on the contingency of association between action and outcome, with the outcome serving as the goal. The motivation behind goal-directed behavior at the individual level is to attain a more desirable outcome. The same logic can be applied to individual learning in a social context. Observing goal-directed behavior of others may motivate the observer to follow the observed action, in case the outcome is expected to be rewarding. An individual is motivated to increase the probability of their own potential reward by correctly following the observed action.

The conceptual connection between goal-directed behavior and learning can be generalized to the concept of *trust*. Trust, as a social capital, has been defined with various connotations. However, the common issue of ’expectation’ emerges as a key determinant of trust across all definitions. An individual’s expectations are tied to his/her predictions. Actions and outcome predictions play pivotal roles in building or destroying interpersonal trust (Rompf, 2015). Thus, the issue of predictability concerns the likelihood of action and outcome occurrence. The constancy of a person’s behavior over time, referred to as consistency, can indicate action predictability. On the other hand, the ability of an individual to successfully execute actions, known as *competence*, determines outcome predictability (Baker, 1987; Rompf, 2015).

The behavior or attitude of a person is deemed trustworthy by the observer if following that specific behavior or attitude proves rewarding to the observer. Hence, learning about the action-outcome contingencies through observational associative learning provides a basis for learning about (emerging trust) and from others (behavioral change). Learning about others may lead the observer to make a decision about forming a trusting relationship with them. This process might result in either trust or distrust towards them, indicating the observer’s future actions (Vecchiato et al., 2014; Wang et al., 2016; Joiner et al., 2017; Eskander et al., 2020; Hertz et al., 2021) In this regard, trust has an undeniable impact on social learning and decision making. Trust can improve the effectiveness of decisions made (Frith & Singer, 2008), based on what has been learnt through observation, here referred to as *observation-based decision making*.

The contributions of different underlying factors, emotion and rationality, in the decision making process is tenable in a trust-based observational learning context (Jones, 1996; Hardin, 2001). The neural activities of distinct emotional and rational structures, proposed by Kahneman (2011)underlie the decision-making process. These activities have been recorded during an individual’s cognitive behaviors in a social context (Smith & DeCoster, 2000; Evans, 2008; Evans & Stanovich, 2013). In the following, we describe major underlying neural structures of the decision-making process in a social context.

The processes described above characterize observational learning and trust-based decision making at the behavioral and social level. However, these processes must ultimately be supported by neural systems that encode value, evaluate observed outcomes, and integrate information relevant for decision making. Understanding how such behavioral phenomena arise therefore requires considering the neural mechanisms that may implement these functions. In the following section, we review candidate cortical systems that have been proposed to support valuation and decision processes involved in observational learning.

#### 1.1.2. The neural basis of social-based decision making

Behavioral processes involved in observational learning and trust-based decision making are thought to rely on neural systems responsible for valuation, prediction, and decision integration. Research in cognitive neuroscience has identified several cortical regions that contribute to these functions, particularly within the prefrontal cortex. These regions have been associated with evaluating observed outcomes, representing expected value, and integrating emotional and cognitive aspects of decision making.

In an earlier study, we presented a neurocomputational model illustrating the involvement of orbitofrontal cortex (OFC) and lateral prefrontal cortex (LPFC) in the human experience-based decision making process (Hassannejad Nazir & Liljenstrom, 2015). OFC and LPFC are also considered to be involved in processing social inputs. However, assessing the integrated social inputs requires yet another neural structure, namely the anterior cingulate cortex (ACC).

OFC has long been known as a structure involved in modulating emotional arousal (Jenison, 2014; Stalnaker et al., 2015). The critical location of this structure profoundly affects its contribution as the recipient of internal efferents from subcortical structures. It contrasts with its role as an integrator of external information from associative sensory structures (i.e., late sensory stimuli).

The strong bidirectional connections of OFC with the amygdala are pivotal to emotional behavior control (Jenison, 2014). The recorded oscillatory activity of OFC represents the expected value of the associated outcome of the afferent stimulus, which is called the *expectancy signal*. OFC puts a high value on the immediate gratification of the associated outcome. Hence, the neural dynamics of this structure are indicative of an individual’s emotional motivation.

The executive functionality of the cortical laminar structure of LPFC was demonstrated by Pribram and others (Pribram, 1987; Mushiake et al., 2006; Figner et al., 2010). The intra-cortical connectivity of the dorsolateral prefrontal cortex serves as an early sensory stimuli processor for subsequent cognitive reasoning (Mushiake et al., 2006; Figner et al., 2010). The oscillatory activity of the LPFC can be represented by the magnitude, time of delivery, and the probability of outcome occurrence of the signal. In contrast to the OFC, the higher valuation of long-term rewards in the LPFC shows the rational aspect of the stimuli. However, both structures pursue goal-directed behaviors (Coutlee & Huettel, 2012; Furuyashiki & Gallagher, 2007). The episodic control of neurons located in the mid-LPFC provides a basis for retrieving maintained episodic memory and directing behavior towards a goal (Asplund et al., 2010; Zald, 2007). Besides, mirror-like neurons were found in LPFC (Simone et al., 2017), indicating that it plays a pivotal role in imitating the observed behaviors of others (Ramsey et al., 2021).

The efferent neurons branching out from LPFC take control of emotional activity by modifying the oscillatory activity of OFC. The power of the afferent neurons to OFC affects the integrated emotional and rational values (Beer et al., 2006; Gray et al., 2002; Hayashi et al., 2013; Hikosaka & Watanabe, 2000; Sokol-Hessner et al., 2012). Therefore, the final individual decision is the result of an integration of the expectancy signal projected from OFC and the rational valuation signal from LPFC.

Burke et al. (2010) illustrate that the oscillatory activities of the ventromedial prefrontal cortex (vmPFC) and dorsolateral prefrontal cortex are determinants of the observational action and outcome prediction errors, respectively. Furthermore, the involvement of OFC in predicting social cues has been recorded by Campbell-Meiklejohn et al. (2012). In this regard, vmPFC/OFC^1^ and LPFC not only underlie emotional/rational individual decision-making, but also serve as bases for observational action-outcome associative learning. The integration of social information projected from these structures occurs in the ACC, which is part of the limbic system. According to documented experiments (Bush et al., 2000; Allman et al., 2001; Kennerley et al., 2011; Hughes & Beer, 2012), the dorsal part of the ACC takes over the rational activities, while the ventral part is responsible for emotional computations. The laminar distributed pattern of the ACC, similar to that of LPFC and OFC, comprises the distribution of interneurons and normal pyramid and spindle cells (Kalus & Senitz, 1996; Palomero-Gallagher et al., 2008).

The contribution of ACC to both rational and emotional aspects of human behavior makes it a hub for cortico-cortical and cortico-limbic connections (Hayashi, 2006). Hence, ACC is regarded as a hub for social valuation. The afferent and efferent neurons connect different neural structures, including LPFC and OFC, to ACC. These connections facilitate the flow of social information within these structures. This attributed function is the result of generating prediction error signals that project from the mentioned structures (Bush et al., 2002; Behrens et al., 2007; Lavin et al., 2013; Anderson et al., 2016). The oscillatory activities of the LPFC and OFC are monitored and updated through the modulatory mechanism of ACC, based on a reinforcement learning process (Medalla & Barbas, 2012;Hill et al., 2016; Joiner et.al. 2017). The schematic diagram illustrating the flow of information in a social context is shown in Figure 1.

**Figure 1.**
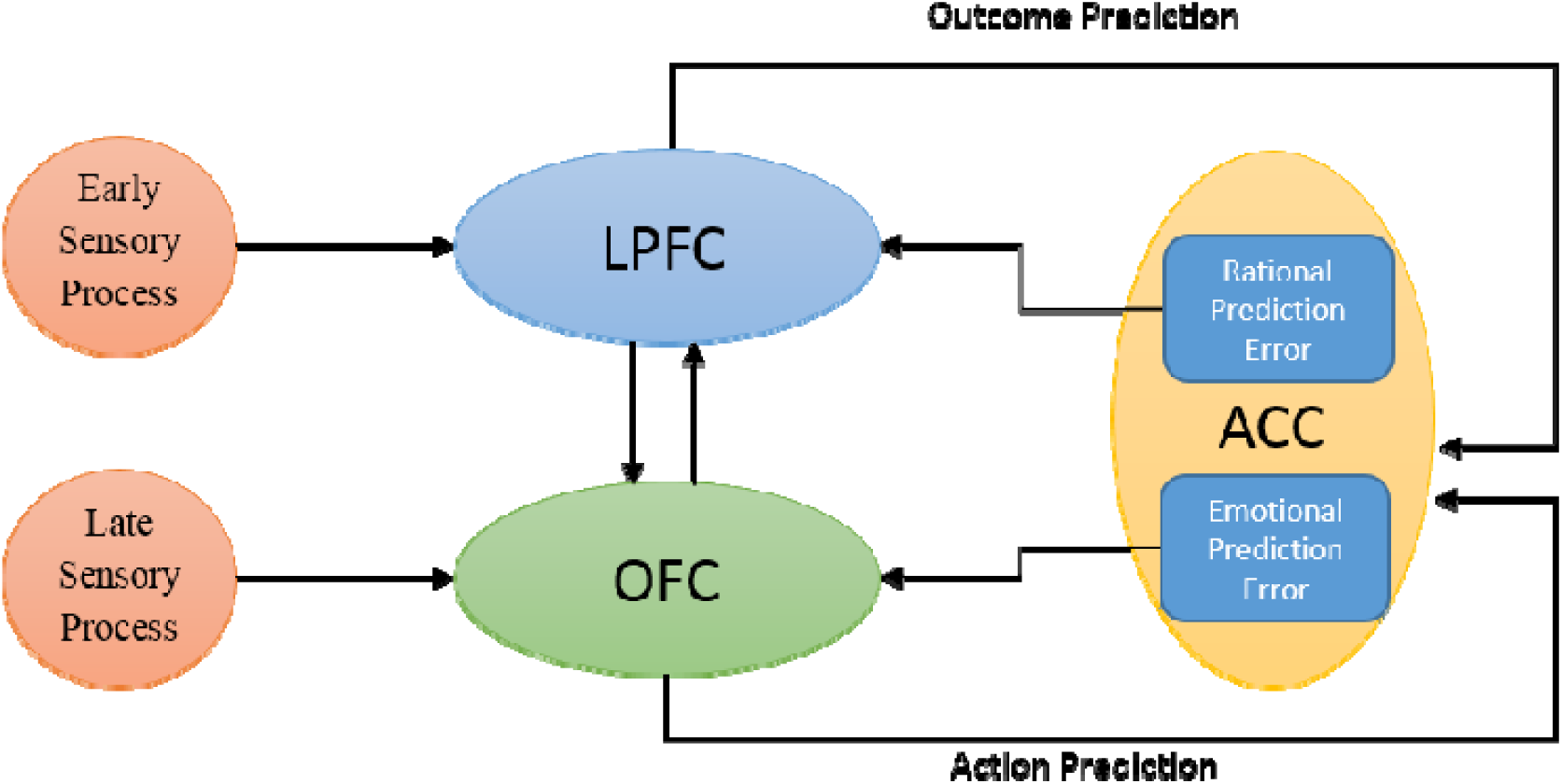
Illustration of the schematic flow of information among three neural structures involved in observational learning. LPFC and OFC are, respectively, subjected to early rational and late emotional sensory stimuli, projecting the processed input to ACC. The differences between the actual and predicted observed action and outcome are signaled as emotional and rational prediction errors, resulting in the update of the oscillatory properties of LPFC and OFC.

Taken together, these findings suggest a correspondence between behavioral processes involved in observational learning and the neural systems that support them. This observation motivates the need for a more explicit framework linking these levels.

#### 1.1.3. Linking social and neural levels

While the above perspectives describe social and neural processes separately, an important step is to clarify how these levels relate to one another within a unified framework. In the present study, social constructs such as trust, observational learning, and decision making are not treated as independent descriptive entities, but are considered to emerge from underlying neural processes involved in valuation, prediction, and integration of information.

In this view, emotional and cognitive aspects of behavior correspond to distinct but interacting neural systems, primarily associated with orbitofrontal and lateral prefrontal regions, respectively. The integration of these processes, together with the evaluation of observed actions and outcomes, provides a basis for understanding how social information can influence individual decisions.

This conceptual linkage serves as the foundation for the neurocomputational model introduced in the following section.

### 1.2. Objectives of this study

Social interactions exert a significant influence on an individual’s attitudes, thereby defining their behaviors and decisions within a social context. While behavioral change may not always stem directly from modifications to the value system, an individual’s attitude could indeed shift. This occurs in response to the motivational value associated with an action.

The level of interpersonal trust is closely linked to societal influences. The behavior of individuals can be influenced or altered when they observe the actions and outcomes of others. This is determined by the level of trust between the observer and the individual whose actions they are witnessing. The concept outlined above forms the conceptual framework of this project. At its core, this study seeks to investigate potential changes in individual behavior when observing an action and its outcome performed by an individual considered skilled (“expert”). This conceptual foundation not only enables an examination of observational learning, but also considers two significant factors: trust and its influence.

In this paper, we explore the reciprocal effects between observational learning and trust. Additionally, we examine how this interaction influences an individual’s emotional and rational assessments, as well as his/her ultimate decision-making.

With our neurocomputational model of the three neural structures, OFC, LPFC, and ACC, we have previously explored the oscillatory neurodynamics activity associated with decision making (Hassannejad Nazir & Liljenström, 2015). Additionally, our model can predict the behavioral pattern of an observer in a trust-dependent observation-based decision-making process. It relies on the oscillatory activities of the three neural structures. The neural pattern properties demonstrate the values of the corresponding decision options and the unidirectional trust relationship between the observer and the observed expert. We can also illustrate the relationship between changes in trust level and changes in emotional/rational attitude towards the decision options. As a measure of the trust level and its influence on observational learning, we consider the predictability of the action and its outcome. The coding of the contingency between action and outcome provides a basis for individuals to learn about and from others in society. The projected action and outcome prediction error signals update and restore the emotional and rational memory about the consistency and competency of the observed expert, respectively. Thus, the observer’s information about the observed expert will be improved. Furthermore, the neural activities involved in associative learning in a social context provide a basis for learning from society. This is achieved through making associations between the observed action and its outcome (Behrens et al., 2008; Heyes, 2012).

Taking into account the variety of decisions we make in our everyday lives, the impact of our decisions related to climate change is of special interest here. A central problem arises concerning how we can change our attitudes and associated behavioral patterns. This is particularly relevant in areas such as travel and transport, which may have a significant impact on climate change. Also in this specific context, our decisions are influenced not only by our own attitudes, but also by the observations we make of others. These observations can range from encounters with strangers on the street to interactions with our neighbors, as well as with more familiar and trusted individuals, such as our family members or even politicians. Hence, we have chosen this example as a case for our neurocomputational model on the neural processes associated with observation-based decision-making processes. In this paper, we explain how emotional and rational values of different means of transport are influenced by trust-dependent observational learning. The characteristics and attitudes of the trustor (observer) and trustee (observed person) make a difference in the quality of the trusted relationship. Consequently, our study is based on specific behavioral and neural assumptions, which will be elaborated upon subsequently.

## 2. Theory and Models

The primary objective of our study is to illustrate how changes in the oscillatory neurodynamics serve as a hallmark of trust-dependent observational learning, impacting the decision-making process. Our goal is to demonstrate the transition of neural activities from heuristic to more rational reasoning, driven by an increase in rational trust.

We demonstrate that the oscillatory activity of LPFC is accountable for measuring the level of rational trust. We evaluate the value of trust by measuring the excitability of neural units, representing an individual’s attitude towards the observed expert.

Our neurocomputational model focuses on trust by evaluating the contingencies of the expert’s action-outcome to determine the impact of these variables on 1) the level of trust and 2) the observer’s decision.

### 2.1. Neurocomputational approach

Our model consists of two parts that focus on the I. individual and II. observation-based decision-making process. The first part focuses on the neural units’ activity, encoding anticipated emotional and rational values of the options. The aim is to study the changes in the observer’s neural behaviors influenced by others. This step is a feed-forward decision process with no immediate feedback. The emotional and rational priorities of the observer, as well as the observation of the expert’s action-outcome association, influence the final decision. It is noteworthy the observer’s predisposition towards the expert’s profession may bias perceptions. This bias potentially leading to positive or negative reevaluation of the expert’s actions, irrespective of their consistency or competence level.

The focus of the second part lies in the neural oscillation involved in vicarious decision-making, considering both the observed action and its subsequent outcome. This process relies on two main factors: 1. Predictions of observed actions and 2. Predictions of observed outcomes, which are generated in the OFC and LPFC, respectively. Discrepancies between the observed (actual) action/outcome and the predicted ones drive changes in the neural properties of the structures associated with both the observed expert and the options chosen by them.

There are different cognitive processes involved in the described process. Mirroring, interactive learning, habitual, and goal-directed valuations are just a few aspects of social learning and decision-making processes. These cognitive processes are underpinned by numerous neural structures. In light of an extensive exploration of this topic, we have focused on modeling two specific neural structures involved in emotional, goal-directed behavior, and decision-making. The orbitofrontal cortex (OFC) primarily handles the encoding of the first two cognitive aspects, while the lateral prefrontal cortex (LPFC) is implicated in the latter two functions. In the social context, the necessity of a coordinating structure that contributes to integrating, monitoring, and controlling socially driven stimuli is evident. Therefore, we consider the activity of the anterior cingulate cortex (ACC). We have developed a neurocomputational model that simulates the decision-making process in a social context based on the prediction and observation of goal-directed behavior. The initial step bears significant resemblance to the neurocomputational model previously developed by Hassannejad Nazir and Liljenström (Hassannejad Nazir & Liljenstrom, 2015) , shown in Figure 2.

**Figure 2.**
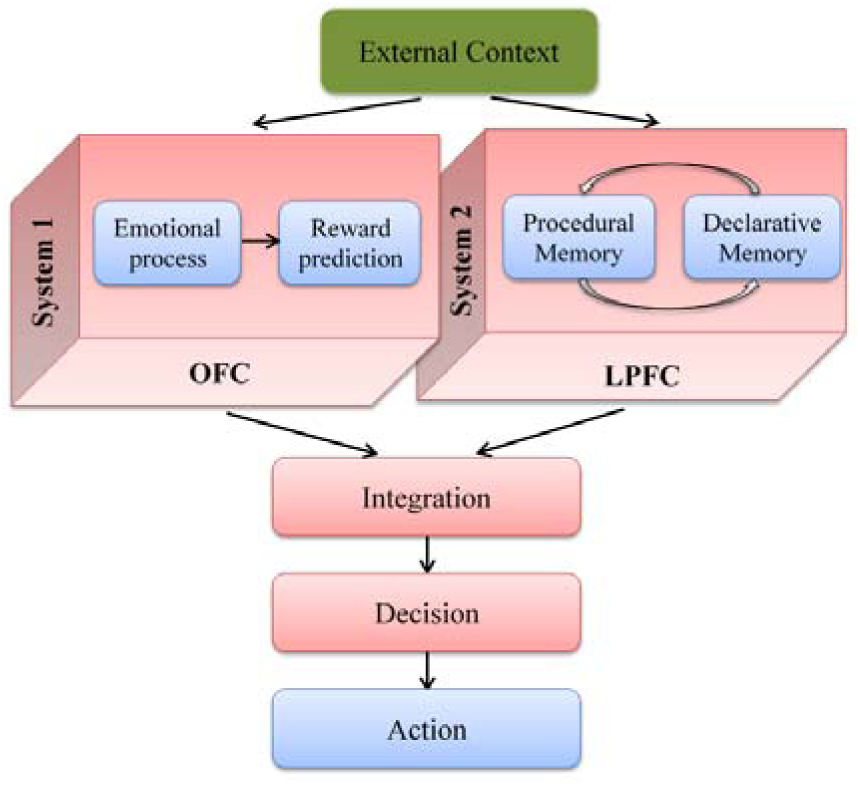
illustrates individual decision-making in the absence of individual experience. Adopted from Hassannejad Nazir and Liljenström (Hassannejad Nazir & Liljenstrom, 2015)

An experience-based decision stems from comparing the anticipated and actual actions encountered by an individual. Conversely, social-based decision-making involves vicarious learning, wherein individuals make decisions based on insights gained from others’ experiences. In our model, the individual decision-making process unfolds in the absence of the individual’s direct experiences. As depicted in Figure 2, the initial part of the individual decision-making process is driven by feedforward neural activities, with the individual’s own experiences and feedback being overlooked.

Figure 3 depicts the schematic demonstration of the second part of the model. Here, the OFC and LPFC are responsible for encoding the anticipated and real emotional and rational values of both the observed expert’s decisions and the vicarious decisions. In LPFC, the neural units signify the rational value of an individual’s decision in the initial stage of the model. Additionally, the involvement of mirror-like neurons in this structure plays a crucial role in assessing the rational value of the observed action-outcome association. The emotional value of the observed action is encoded by OFC, representing its desirability. The orbitofrontal and lateral neurons transmit the anticipated (3.a) and actual (3.b) values (both emotional and rational) of the expert’s action-outcome association to ACC. The integration of these signals in ACC generates errors in observational action and outcome prediction. The predicted and actual values of the expert’s decisions are influenced by the individual’s emotional-rational experiences, priorities, and prior bias towards the expert.

**Figure 3.**
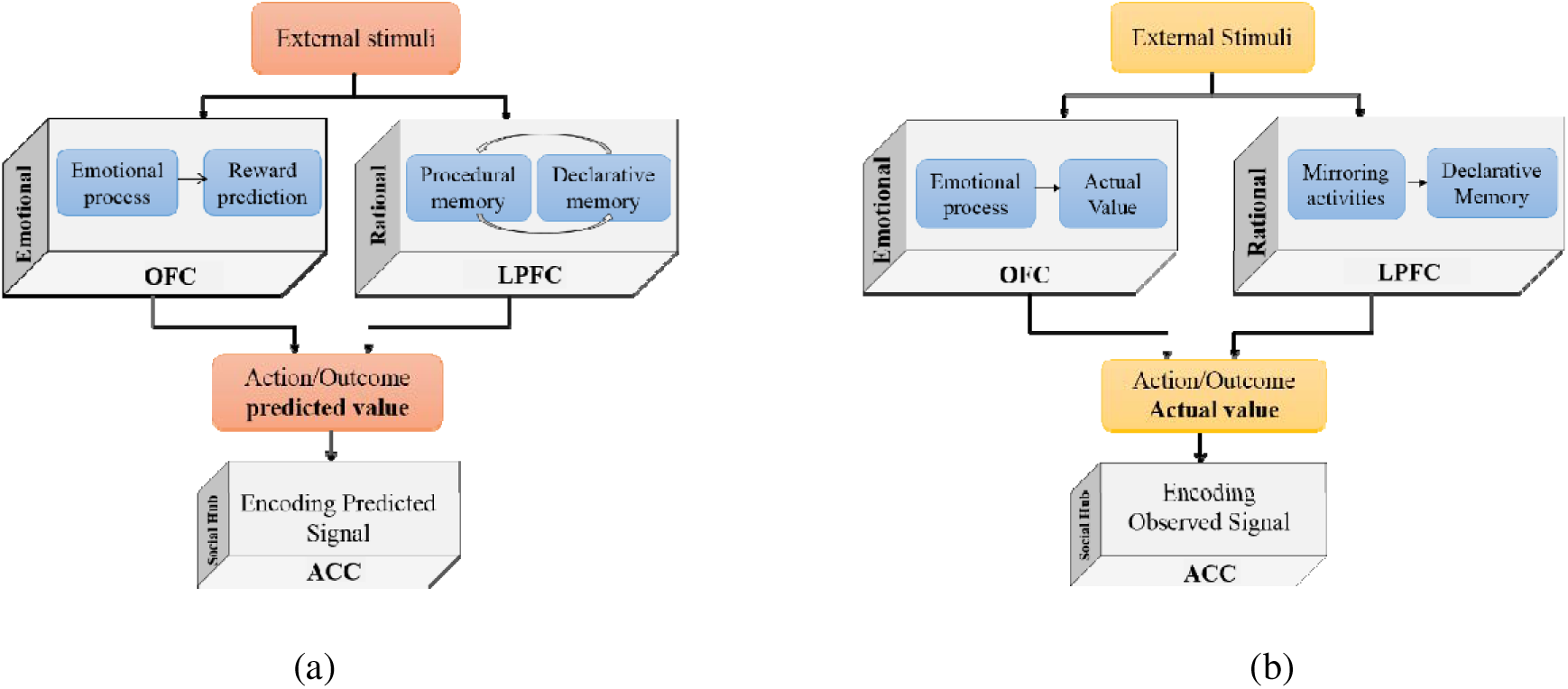
illustrates (a) the schematic demonstration of observational action/outcome prediction and (b) action/outcome observation from a neural viewpoint. The emotional and rational value of the observed action is computed vicariously in the OFC and LPFC, respectively. The final value is the result of signal integration, which is projected into the ACC for further processing.

The encoded observational action and outcome prediction errors in the ACC are projected to the OFC and LPFC to update the properties of neural patterns associated with the observed action and the expert, as depicted in Figure. 4.

**Figure 4.**
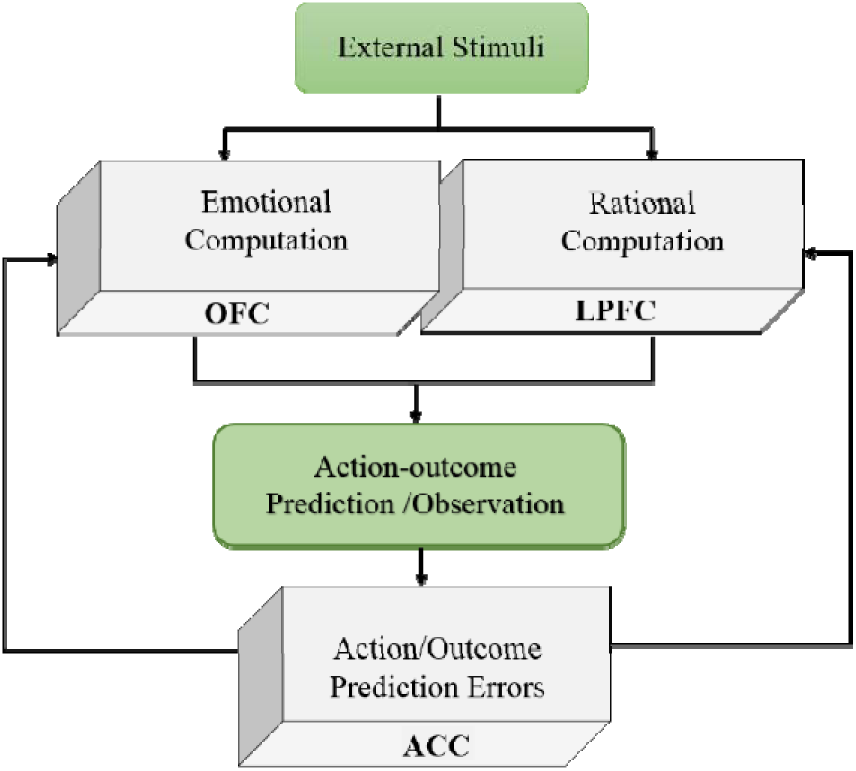
Integration of predicted and actual values. The difference between the values of predicted and actual afferent signals from the OFC and LPFC generates an error-related negativity signal. This signal is then sent back to the emotional and rational structures to update the neural variables such as learning rate, oscillatory properties, and excitability of neural units.

#### 2.1.1. Neural structures

Despite the variations observed in the structures of the OFC, LPFC, and ACC, all these structures are organized in layers. Taking this into account, Hassannejad Nazir and Liljenström (Hassannejad Nazir & Liljenstrom, 2015) modeled the neurodynamics of the OFC and LPFC based on a three-layered neural model developed by Liljenström (Liljenström, 1991).

Considering the neuroanatomy of the ACC, the structure is designed similarly to the OFC and LPFC to avoid the complexity caused by the diversity of cellular composition in the ACC. The upper and lower levels of the three-layered structure comprise inhibitory neurons corresponding to layers III and Vb of the ACC, while the middle layer represents the high density of excitatory cells in layer V of the ACC with excitatory neural units. Hence, the three-layered structure is common among all three structures. This structure is suggested based on the attractor neural network, which provides a basis to generate the oscillatory activities of neural patterns. The modeled activity of cell assemblies represents the processes of memory formation, retrieval, valuation, and prediction. The neural interactions between the middle excitatory neurons and the two feedforward and feedback inhibitory layers, stimulated by external stimuli, are illustrated in Figure 5. Although the structures of these neural parts are notably simplified, the accuracy of their performance has been considered in the developed model.

**Figure 5.**
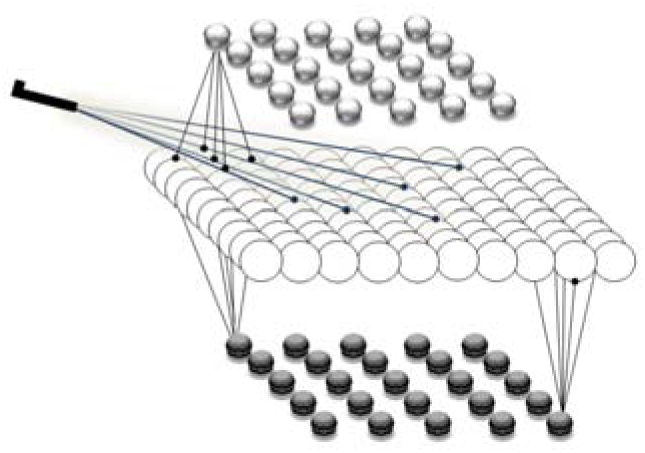
Schematic illustration of a simplified three-layered neural structure consisting of excitatory and inhibitory neural units. The excitatory layer located in the middle is composed of 100 neural units, with two feedforward and feedback inhibitory layers at the top and bottom, respectively, each containing 25 neuronal units.

The oscillatory activities of the neural units have been computed as follows:

(1)

while *u_i_* represents the internal states of the neural unit stimulated by the external input, *I(t)*. In the model, the matrix *W* represents the connectivity structure between neural populations. Each element of this matrix denotes the strength of the connection from unit *j* to unit *i*, with a conduction delay, *δ_ij_*, and the membrane time constant, *τ_i_*, play role in measuring the rate of neural activity changes.

The input-output function, *g_i_*(*u_i_*) measured by Freeman is based on the gain parameter, *Q_i_*, denoting the level of arousal, or the excitability of units to form neural patters(Freeman, 1979). C is the constant value.

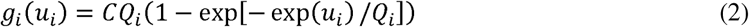

Here, the learning process follows the Hebbian theory (Hebb, 1952) :

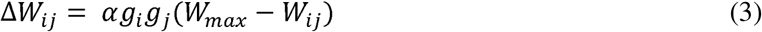

where α, is the learning rate. The parameter *W_max_* represents the maximum allowable connection strength between neural units in the model. It constrains the range of synaptic weights and ensures that the strength of interactions between neural populations remains within biologically plausible limits. The impact of trust on social processes has been considered in the literature. Social learning is an important constitute of social processes, also influenced by trust. Hence, to consider the correlation between neural learning and trust, we modified the Hebbian learning equation as follows:

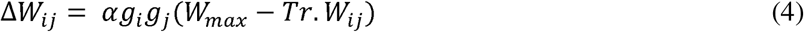

where *Tr* is a variable, proportional to the trust level. The more consistent and competent the observed expert is, the higher the *Tr* value will be. Initially, *Tr* equals to 1, following the general Hebbian learning. The variable *Tr* takes a higher value when the observer’s trust in the expert changes. An increase in *Tr* increases the rate of trust learning and memory consolidation. Therefore, the neural oscillatory activities of cell assemblies, signal energy and intensity, associated with options change during observational learning.

We consider the value of each decision option, *V*(*opt*), to be proportional to the weight strength, *E_opt_*, and energy of the corresponding cell assembly, *W_opt_*. The energy of the cell assembly is based on the signal’s frequency, *E_opt_*, and amplitude, *A_opt_*.:

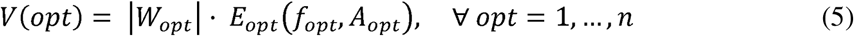

#### 2.1.2. Neural aspect of probability theory

As discussed above, the observational action and outcome predictions are two indivisible aspects of the social-based decision-making process. The prediction of others’ actions and the subsequent outcomes evolves out of the past observed actions–outcomes associations. The likelihood of occurrence of an action and outcome are the measure of the action and outcome prediction.

Understanding how the brain encodes probability and the neural representation of probability are crucial topics in neuroscience. Prominently, it is presumed that the neural activities in an uncertain environment can be modeled by Bayesian theory (Doya, 2007; Knill & Pouget, 2004; Rich et.al., 2015; Schultz & Dickinson, 2000).

The Bayesian theorem is a method that calculates the probability of an event occurring in a single time slot based solely on the current state. On the contrary, Dynamic Bayesian Probability (DBP) (Ghahramani, 1998; Lin & Mitchell, 1992) is a suitable method, and it is essential to consider the sequence of actions. The following equation represents the probability of the occurrence of an action at time n+1, *X*_*n*+1_, given the past actions based on DBP.

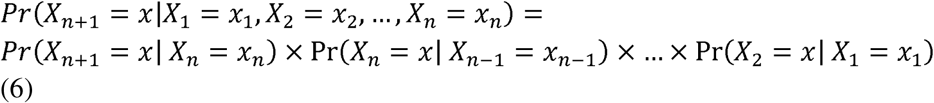

This method can be applied for active learning problems while environmental changes are important. Given the history-dependent nature of decision-making, the likelihood of the expert making a decision hinges on the associative prediction of past actions and outcomes. The overall approach and attitude of the expert can be inferred through a sequential dependent probability distribution. Therefore, DBP is applied here to regulate the neural oscillatory activities during observations. The strength of the predictive signals (i.e., mean peak value and frequency) is computed based on the probabilities of observed actions and outcomes.

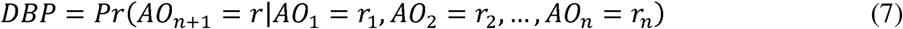

*AO* represents the action-outcome association, while *r_n_* denotes the desired rewarding outcome at time *n*. In this study, the neural coding of probability is measured based on the neural excitability, strength of neural unit connections, and associative connections’ strength. Additionally, we apply the computed probability, *DBP*, to measure the properties of neuronal units encoding uncertainty. Signal intensity is determined by signal energy (*E_s_*), dynamic probability (*DBP*), and motivation to follow the observed individual (*DBP*), i.e., trust in the observed individual.’

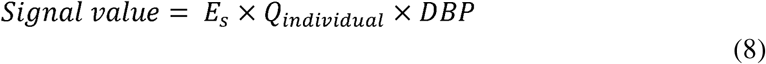

The signal value represents the magnitude of the prediction signals, indicating the action and outcome predictability of the expert, as well as the probability that the observer follows the expert.

## 3. Simulation and Results

### 3.1. Assumptions

In this study, the conceptual model is based on the recorded observational action and outcome prediction error in the prefrontal cortex areas. The conceptual models and the applied cortical model are illustrated in Figures (1), (4) and (5). The influences of social information on adaptive heuristic and rationality of human decision-making are simulated and analyzed with respect to some initial behavioral and neural assumptions.

The observation-based decision-making process is influenced by the characteristics of both the observer and the observed individual. To investigate the transition from emotional to rational reasoning, we assume that the observer initially exhibits higher emotional propensity compared to rational tendencies. Therefore, the intensity of emotional signals, motivation (Q), and excitability of neural populations are designed to be higher than the rational counterparts. Different Q values representing varying levels of emotional and rational motivations are presented in Table 1, along with the values of variables in equations (1-4).

**Table 1.**
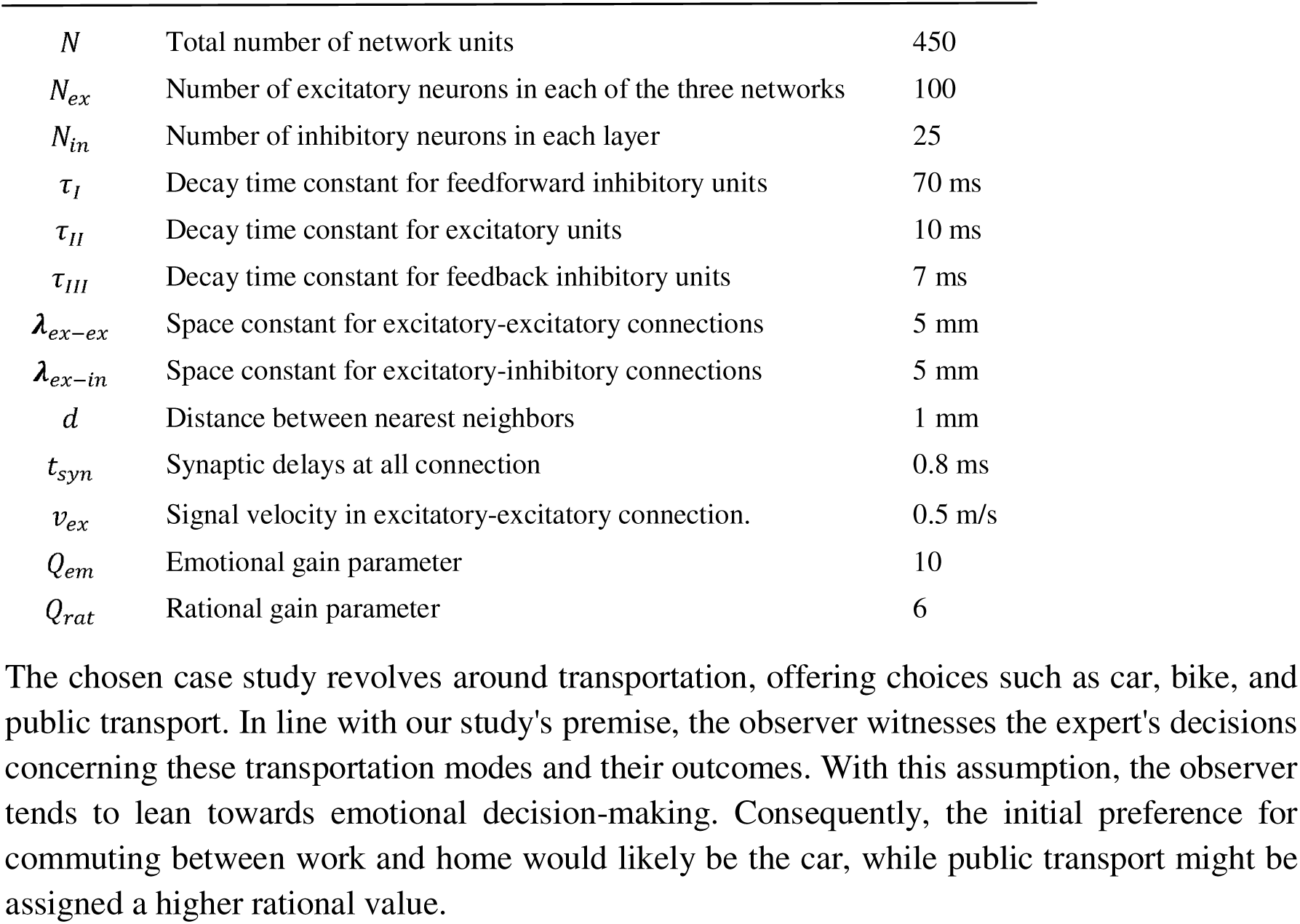
The parameters chosen for the simulation of the three-layered networks in each of the three structures, ACC, OFC and LPFC. Many of these numbers are within physiological realistic ranges.

|  |  |  |
| --- | --- | --- |
| $N$ | Total number of network units | 450 |
| $N_{ex}$ | Number of excitatory neurons in each of the three networks | 100 |
| $N_{in}$ | Number of inhibitory neurons in each layer | 25 |
| $\tau_I$ | Decay time constant for feedforward inhibitory units | 70 ms |
| $\tau_{II}$ | Decay time constant for excitatory units | 10 ms |
| $\tau_{III}$ | Decay time constant for feedback inhibitory units | 7 ms |
| $\lambda_{ex-ex}$ | Space constant for excitatory-excitatory connections | 5 mm |
| $\lambda_{ex-in}$ | Space constant for excitatory-inhibitory connections | 5 mm |
| $d$ | Distance between nearest neighbors | 1 mm |
| $t_{syn}$ | Synaptic delays at all connection | 0.8 ms |
| $v_{ex}$ | Signal velocity in excitatory-excitatory connection. | 0.5 m/s |
| $Q_{em}$ | Emotional gain parameter | 10 |
| $Q_{rat}$ | Rational gain parameter | 6 |

In addition to the significance of the observer’s attributes in decision-making, the predictability of the expert is crucial. In this study, we investigate the impact of an expert with a high level of predictability, measured in relation to the assumed profession of the expert (e.g. a professor in climatology). The trust level in this expert is set at a rather high level (i.e. 70%), with the strength of neural connections representing initial trust levels. Trust undoubtedly arises from both emotional and rational foundations, but for simplicity, we focus on the rational aspect.

The chosen case study revolves around transportation, offering choices such as car, bike, and public transport. In line with our study’s premise, the observer witnesses the expert’s decisions concerning these transportation modes and their outcomes. With this assumption, the observer tends to lean towards emotional decision-making. Consequently, the initial preference for commuting between work and home would likely be the car, while public transport might be assigned a higher rational value.

### 3.2. Neural representation of trust

The excitability of the neural units depends on the strength of the neural connections, which is influenced by intra-neural stimulation. Strong neural connections are capable of creating more accurate and coordinated neural patterns, resulting in simultaneous oscillations with high energy. The influence of observational learning on an individual’s decision-making process manifests in the neural connection weights and neural excitability. The oscillatory properties of neural patterns associated with the expert reflect the level of trust. Figure 6 illustrates the changes in LPFC’s oscillatory activities during observational learning. EEG-like data demonstrate varying levels of trust based on the observer’s learning about the expert. The neural connection weights and motivation (Q) of the corresponding pattern determine the excitability of neural units, with different levels of trust emerging from different Q values. The neural arousal variable, Q, indicates the observer’s motivation to follow the expert’s actions. Thus, as Q increases, so does the level of trust. Additionally, the dominant frequency of neural activity reflects signal energy. A higher dominant frequency in a neural pattern signifies greater trust. For instance, in Figure 6.a, the dominant frequency of the corresponding neural pattern for expert 1 (trust level) is 50 Hz, with a Q value of 6. Increasing the Q value to 8 and 9 results in dominant frequencies of 60 and 79, respectively, indicating higher levels of trust.

**Figure 6.**
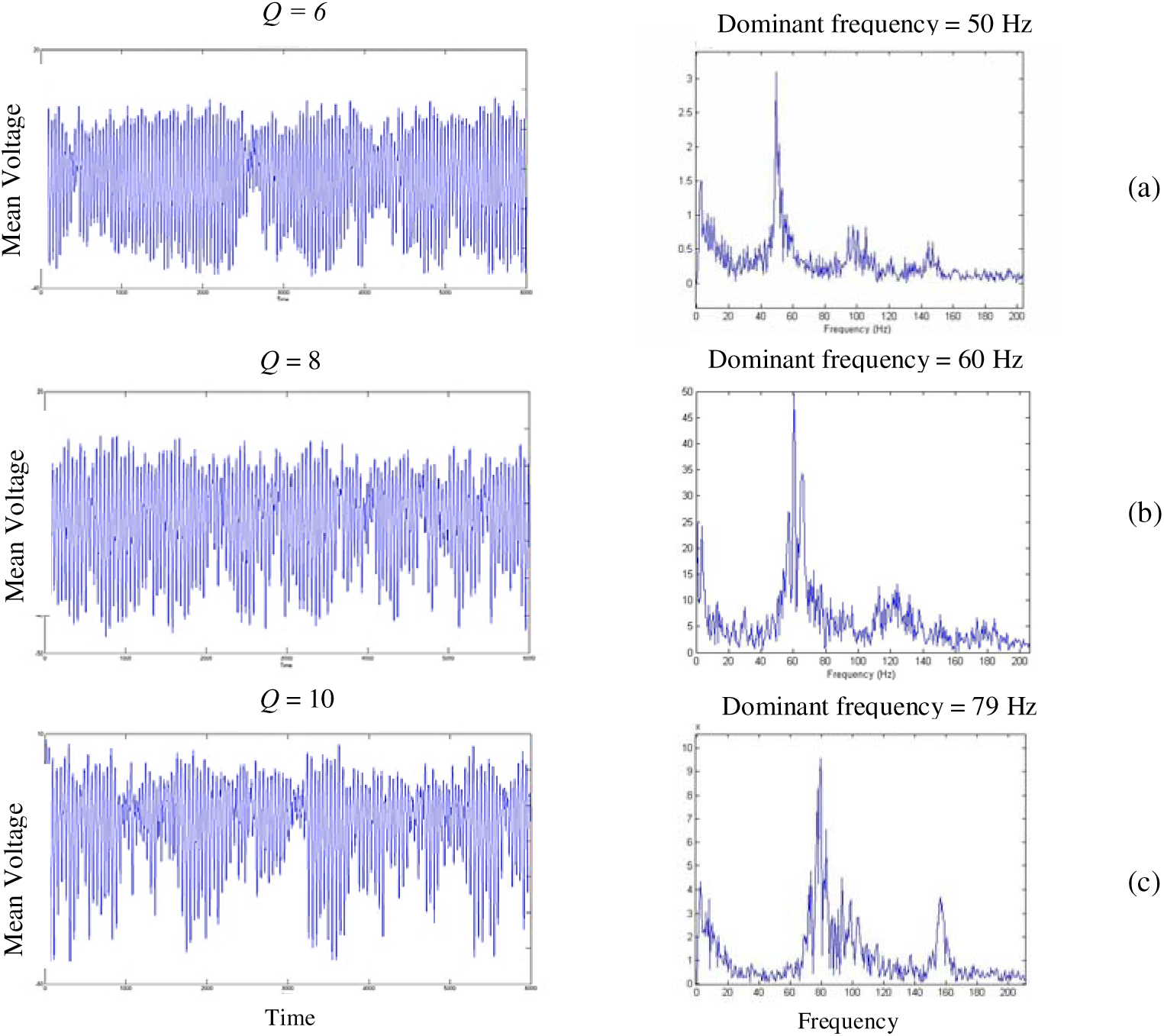
oscillatory activities of LPFC’s neural pattern corresponding to the public transport. The increase in trust brings about the higher rational neural excitability and the higher rational motivation. The higher the motivation becomes, the more regular with higher frequency the neural activity will be.

Next, we explore the behavioral change of the observer from two different perspectives: 1) the impact of observational learning on trust, and 2) the impact of trust on observational learning.

### 3.3. Impact of observational learning on trust

The predictive signal representing the observer’s decision is proportional to the contingency of the action-outcome association. An expert with a high level of predictability (both in action and outcome) positively influences the rational trust of the observer. The level of trust in the expert is measured based on the oscillatory activities of the LPFC’s neural patterns. According to Eq. 8, a high value of DBP results in high signal energy and an excitable neural pattern. The excitability of the corresponding neural pattern is computed as a function of motivation, Q, and the neural connection weights. Changes in the neural properties demonstrate the learning process leading to behavioral changes in the observer.

Figure 7 shows the trust learning curve. The changes in the connection weights during the observational learning process result from changes in the Q value, representing different levels of trust in the expert. The rate of changes in the neural connection weights decreases during the observation of the expert, and the connections will eventually saturate, determining the stability of the observer’s trust in the expert. The increase in the strength of neural connections demonstrates an increase in the excitability of the cell assembly associated with the observed individual.

**Figure 7.**
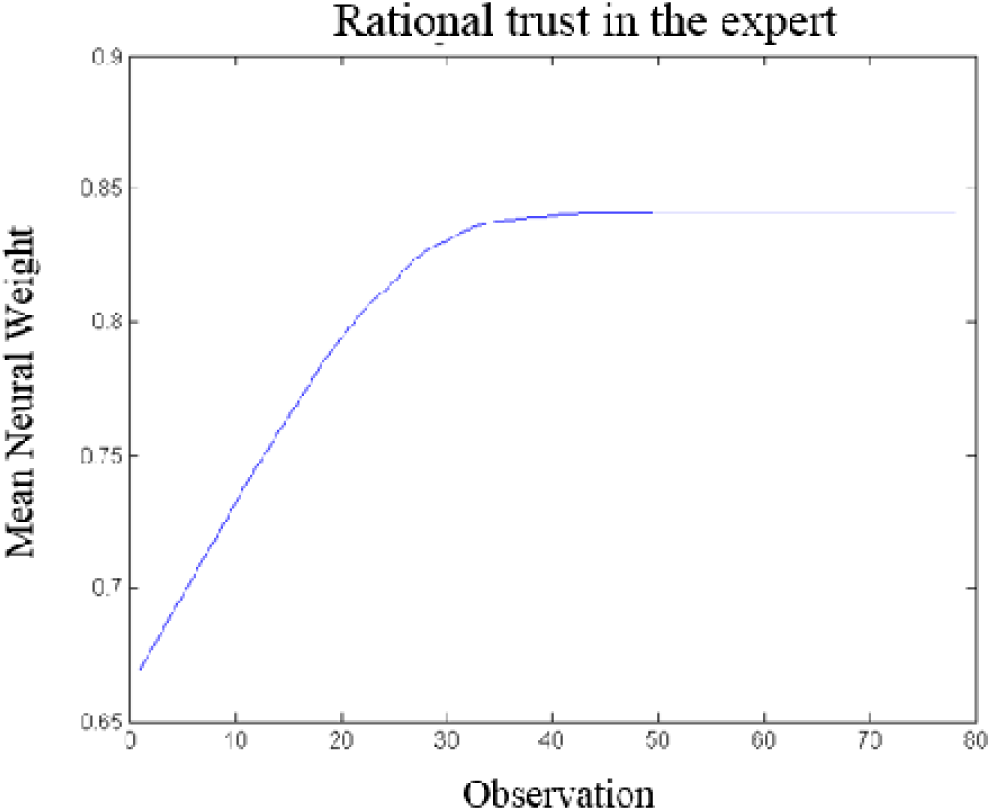
Illustration of the trust changes through observing the action-outcome of an expert. After 40 iterations, the level of trust reaches the steady state triangle sign).

An increase in the trust level results in an increase in the rational and emotional values of the observed action, as described below.

### 3.4. Impact of trust on observational learning

The attitude of the observer towards public transport (the expert’s action) is influenced by trust in the expert. In this part, we study the changes in the observer’s emotional and rational valuation according to changes in trust.

The strength of the rational signals projected by the LPFC’s cell assembly, corresponding to the expert, reflects the level of rational trust. The observer’s stronger inclination towards choosing the car suggests that the expectancy signal of the corresponding OFC neural pattern outweighs that of the neural pattern associated with public transport. However, the emotional neural pattern linked to public transport exhibits lower excitability and weaker connection weights compared to its rational neural counterparts. As depicted in Figure 8, the observer initially demonstrates a higher rational inclination towards selecting public transport than its emotional value. The enhancement in rational trust, as depicted in Figure 7, positively impacts the observer’s valuation process. This leads to an increased rational value attributed to the expert’s action (i.e., public transport) from the observer’s perspective.

**Figure 8.**
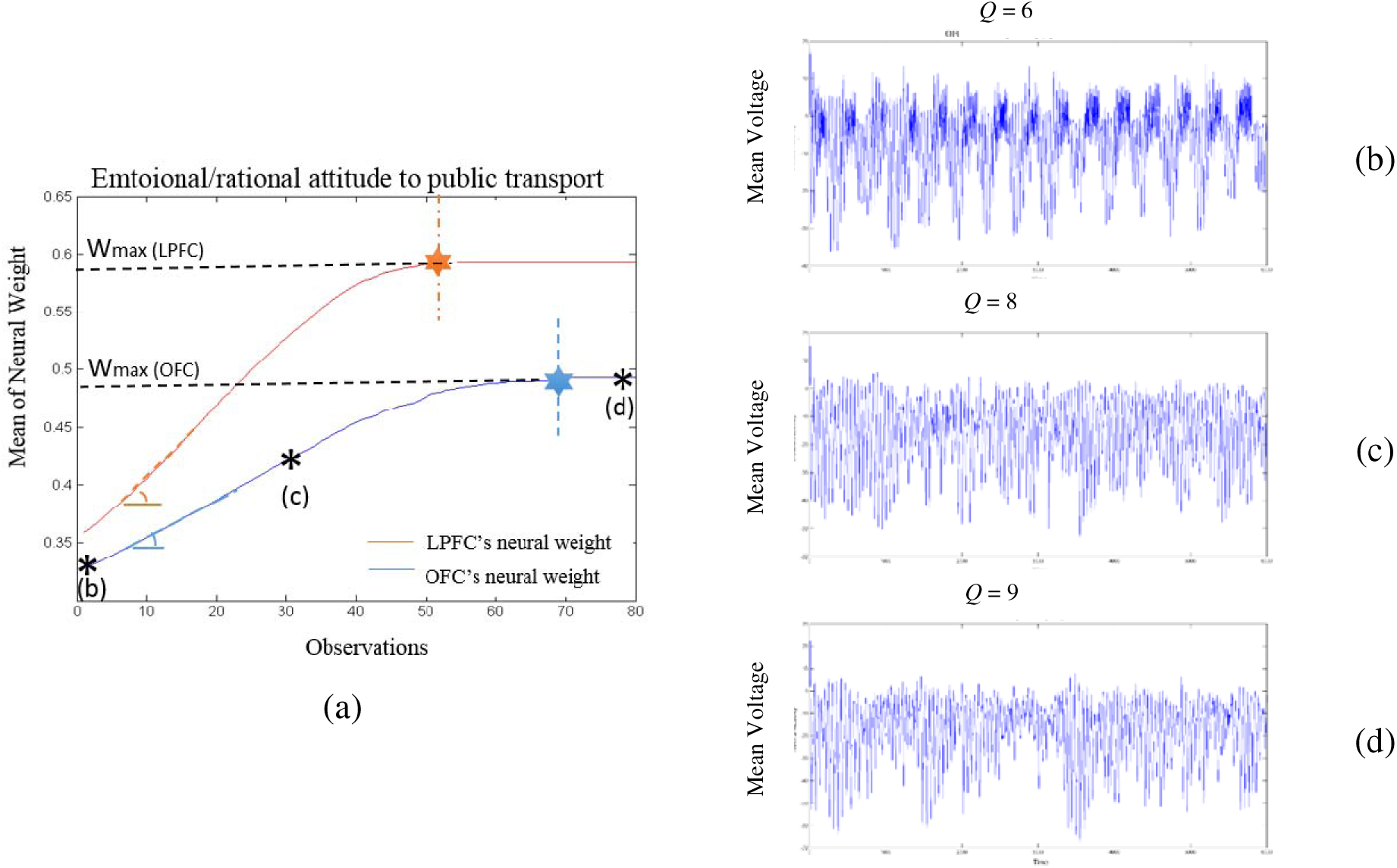
Illustration of the emotional and rational changes in the neural connection weights corresponding to the public transport pattern (a).As an assumption, the emotional tendency to select public transport is less than the rational propensity. Observing an expert selecting the car with a successful outcome augments the rational valuation of public transport, albeit with a lower rate, and diminishes the emotional value. The oscillatory activities of OFC’s neural units associated with public transport are illustrated in Figures (b), (c), and (d). The emotional Q values increase influenced by LPFC’s modifications.

The increased oscillatory activity of the LPFC’s neural pattern modifies the oscillatory rhythms of OFC’s neural pattern encoding public transport. Hence, the strength of OFC’s cell assembly corresponding to public transport increases, but at a lower rate compared to the increase in the LPFC’s neural pattern. Figures 8.b, 8.c, and 8.d illustrate the changes in the oscillatory activity of the LPFC’s cell assembly associated with public transport during the observational learning process. The in the first frame is 6, and the neural weight connection will stabilize when the Q value reaches 9. Thus, the rational neural connection weights become saturated (Figure.8.a. orange dotted line) before the emotional neural pattern reaches its maximum neural strength (Figure 8.a. blue dotted line). Additionally, the maximum weight of the LPFC’s neural pattern is considerably greater than the saturation level of the OFC’s corresponding pattern.

In the following section, the properties of the emotional and rational cell assemblies associated with public transport during the observation of the expert’s actions and outcomes are measured. The excitabilities of the OFC’s and LPFC’s cell assemblies before the decision maker starts observing the action and outcome of the expert are as follows:

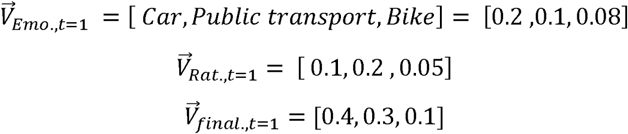

Here, we assume the decision maker is a more emotional than rational person. Initially, she is more inclined to choose the car. However, her rational preference is to take public transport. In line with our assumption, the initial emotional value of public transport is less than its rational value. Hence, the mean neural weight of the emotional pattern associated with public transport is less than the rational neural weight of this option. The strength of the cell assembly corresponding to public transport after observing the action-outcome association of the expert is measured as follows:

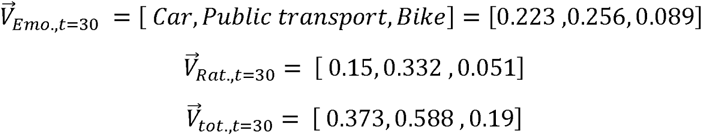

Making a comparison between Figures 7 and 8, the trust learning rate differs from the rate of changes in emotional/rational reasoning. The results illustrate that the learning rate of trust in expert 1 is higher than the rate of emotional/rational learning about the expert’s action (i.e., public transport). Therefore, the required time for attitude change (towards public transport) is longer (almost twice as long) than the required time to form the attitude towards the expert.

## 4. Discussion

In this paper, we have presented our neurocomputational model, exploring the interplay between trust and observational learning, as well as the impact of social influence on an individual’s decisions. Our model is an attempt to demonstrate possible interactions between three cortical structures: OFC, LPFC, and ACC in a social-based decision-making context. The involvement of OFC and LPFC in individual and social cognitive processes, as well as their interaction with ACC as a social hub, provides the framework for our model. The emotional and rational analyses of personal attitude and social information are the major contributions of OFC and LPFC, respectively, in this model. The conceptual grounding of this model is based on observational action and outcome prediction error signals recorded (as simulated) in OFC and LPFC, respectively. At the neural level, learning is defined based on the difference between the predicted and actual values in individual and social contexts. Hence, in this model, we take advantage of the observational prediction error signals to provide a basis for the observer to: 1) learn about and from the observed expert, and 2) modify individual decisions and probably attitudes in the long term. Concerning the gathered knowledge about the expert, the observer makes an association between the observed action-outcome pair and the expert, which results in an increase/decrease of trust.

This model predicts the decision pattern of a new observer in the observation-based decision-making process. It also gains an understanding of the interplay between trust and observational learning. The model predictions are based on the analyses of cell assembly activities (signal intensities and neural excitability).

The illustrated results can be analyzed from both neural and behavioral perspectives. As shown in Figure 6, there is a direct relationship between the properties of cell assemblies and the level of trust. A higher level of trust is reflected in stronger neural connection weights and higher neural excitability, resulting in less chaotic behavior. The simulated data illustrates how the oscillatory activities of neural patterns project the expectancy signal of option values. The higher the motivation becomes, the more regular the oscillatory activity will be.

Figure 8 depicts the oscillatory neurodynamics of both rational and emotional responses concerning the predictability of actions and outcomes resulting from the expert’s actions. Based on the simulation results, changes in emotional and rational values do not occur simultaneously. Rational value adjustments, influenced by rational trust, manifest at a faster rate compared to emotional changes. The stabilization of rational oscillatory activity is observed sooner than that of the emotional counterpart. Despite the generally faster nature of emotional reasoning, as noted by Kahneman (2011), we conclude that alterations in emotional preferences or attitudes require more time than changes in rational attitudes.

In summary, observing the actions and outcomes of others serves as a robust basis for both learning from and about them. Individual learning about others, driven by the predictability of their action-outcome associations, underpins the development of trust. Trust, along with the knowledge acquired from others, significantly shapes both emotional and rational valuation processes. Consequently, observational learning and trust emerge as two pivotal social variables profoundly influencing individuals’ behaviors and attitudes.

## Footnotes

1 Despite the differences between vmPFC and OFC, some structural overlaps and functional similarities between these two structures make it possible for us to replace vmPFC with OFC in our social-based decision making. The similarities between the neural arrangements in terms of Brodmann area as well as to some extent similar cortico-cortical connections have been observed in these structures. In addition, both structures underlie many similar cognitive processes such as emotional processing and decision making.

